# Neural correlates of conscious visual perception lie outside the visual system: evidence from the double-drift illusion

**DOI:** 10.1101/660597

**Authors:** Sirui Liu, Qing Yu, Peter U. Tse, Patrick Cavanagh

## Abstract

When perception differs from the physical stimulus, as it does for visual illusions and binocular rivalry, the opportunity arises to localize where perception emerges in the visual processing hierarchy. Representations prior to that stage differ from the eventual conscious percept even though they provide input to it. Here we investigate where and how a remarkable misperception of position emerges in the brain. This “double-drift” illusion causes a dramatic mismatch between retinal and perceived location, producing a perceived path that can differ from its physical path by 45° or more [1]. The deviations in the perceived trajectory can accumulate over at least a second [1] whereas other motion-induced position shifts accumulate over only 80 to 100 ms before saturating [2]. Using fMRI and multivariate pattern analysis, we find that the illusory path does not share activity patterns with a matched physical path in any early visual areas. In contrast, a whole-brain searchlight analysis reveals a shared representation in more anterior regions of the brain. These higher-order areas would have the longer time constants required to accumulate the small moment-to-moment position offsets that presumably originate in early visual cortices, and then transform these sensory inputs into a final conscious percept. The dissociation between perception and the activity in early sensory cortex suggests that perceived position does not emerge in what is traditionally regarded as the visual system but emerges instead at a much higher level.

## Introduction

The representation of location is determined by an object’s current retinal location in combination with several other sources of information, such as head and eye directions [3-6], eye movement plans [7], and the object’s own motion [8]. Studies have shown that our visual system can predict the current location of a moving target by taking into account its velocity and the neural delays between the retina and the cortex [9]. This predictive position shift, extrapolating the target ahead along its motion path, was proposed to underlie several motion-induced position shifts in which an object’s location appears to be shifted by surrounding motion signals or by its own motion [8,10-13].

The goal of the present study is to use predictive position shifts to investigate where the representation of perceived position emerges in the processing hierarchy. We used a probe that induces a remarkably large motion-induced position shift, namely, the double-drift illusion [1,14-16]. Compared with other well-known motion-induced position shifts, this stimulus reveals an integration of motion signals over a second or more, leading to dramatic shifts in perceived position that can deviate from the physical motion trajectory by many visual degrees (**Figure 1** and **Movie S1**). With such a long integration period, it is unlikely that early visual areas with their short integration time constants are responsible for the accumulation of position errors underlying this illusion. Thus, the double-drift stimulus presents the opportunity to explore where in the visual processing hierarchy position information transitions from retinally-based, bottom-up encoding, to high-level, motion-influenced perceptual representations associated with visual conscious experience. Specifically, if the patterns of neural activity that encode perception can be distinguished from those driven by the physical stimulus, we can identify the cortical areas where the percept first arises using multivariate pattern analysis (MVPA) on fMRI signals.

**Figure 1.**
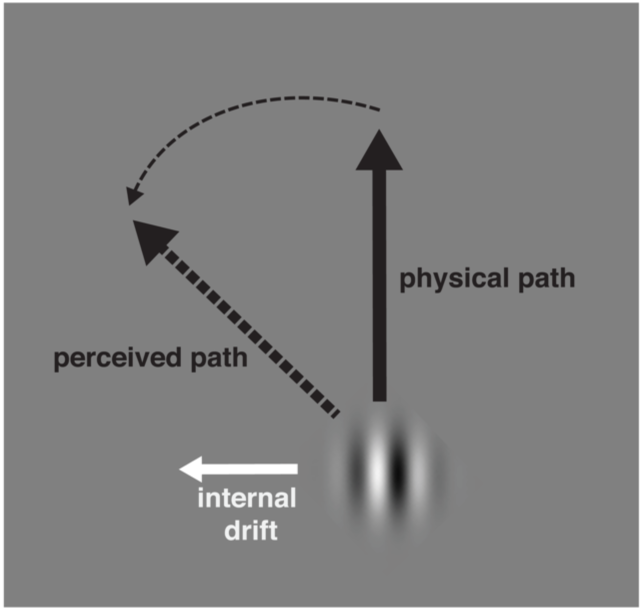
Double-drift stimulus. A Gabor patch with vertical physical motion path can be perceived to be moving obliquely if its internal texture is drifting orthogonally to its physical path. See also Movie S1.

The present study investigated where and how the perceived position of the double-drift illusion is encoded in the brain using fMRI and MVPA. We found that the illusory motion paths of two double-drift stimuli with identical retinal but different perceived paths can be decoded from multiple brain regions, but the nature of the representation differs in these brain areas: activation patterns for the illusory motion paths were decodable in V2 and V3 but not in V1 or MT+. However, cross-classification between the double-drift and control stimuli with matched physical motion paths showed no evidence that the pattern of response to the illusory paths in these areas was related to the response to their matched physical paths. In contrast, a whole-brain cross-classification searchlight analysis revealed that activations in more anterior parts of the brain share a common position encoding for the illusory and matched physical path of the double-drift stimulus with no such shared representation observed in early visual areas. That is, only in higher-order brain areas did the neural coding reflect the similarity in perception of the illusory and matched physical paths. Thus, our results indicate that different cortical regions are involved in representing different properties of the double-drift stimulus, with the early retinotopic visual areas V2 and V3 possibly generating the local direction deviations driven by motion signals integrated over short durations and the higher-order regions possibly accumulating and storing position displacements based on extrapolations of those integrated motion directions to represent the long-lasting perceived motion path that can deviate from its physical path by 45° or more.

## Results

### Perceived path orientation of the double-drift stimulus deviates largely from its physical path orientation

We first conducted a behavioral task to measure the size of perceived position shift of the double-drift stimulus for each participant (see **Figure 2A** and **STAR Methods** for details). As expected from previous literature, the perceived path orientation of the double-drift stimulus was significantly different from that of the control stimulus that lacked internal motion (perceived rightward tilt: *p* < .001, Cohen’s *d* = 13.42; perceived leftward tilt: *p* < .001, Cohen’s *d* = 14.89). Specifically, the perceived path orientation was biased toward its internal drift direction, suggesting that there was a consistent motion-induced position shift of our double-drift stimuli across all subjects (average illusion size = 47.55°). There was no significant difference in the absolute amount of perceived direction shift between the two internal drift conditions (i.e. leftward vs. rightward tilt) of the double-drift stimulus (*p* = .67, Cohen’s *d* = .23).

**Figure 2.**
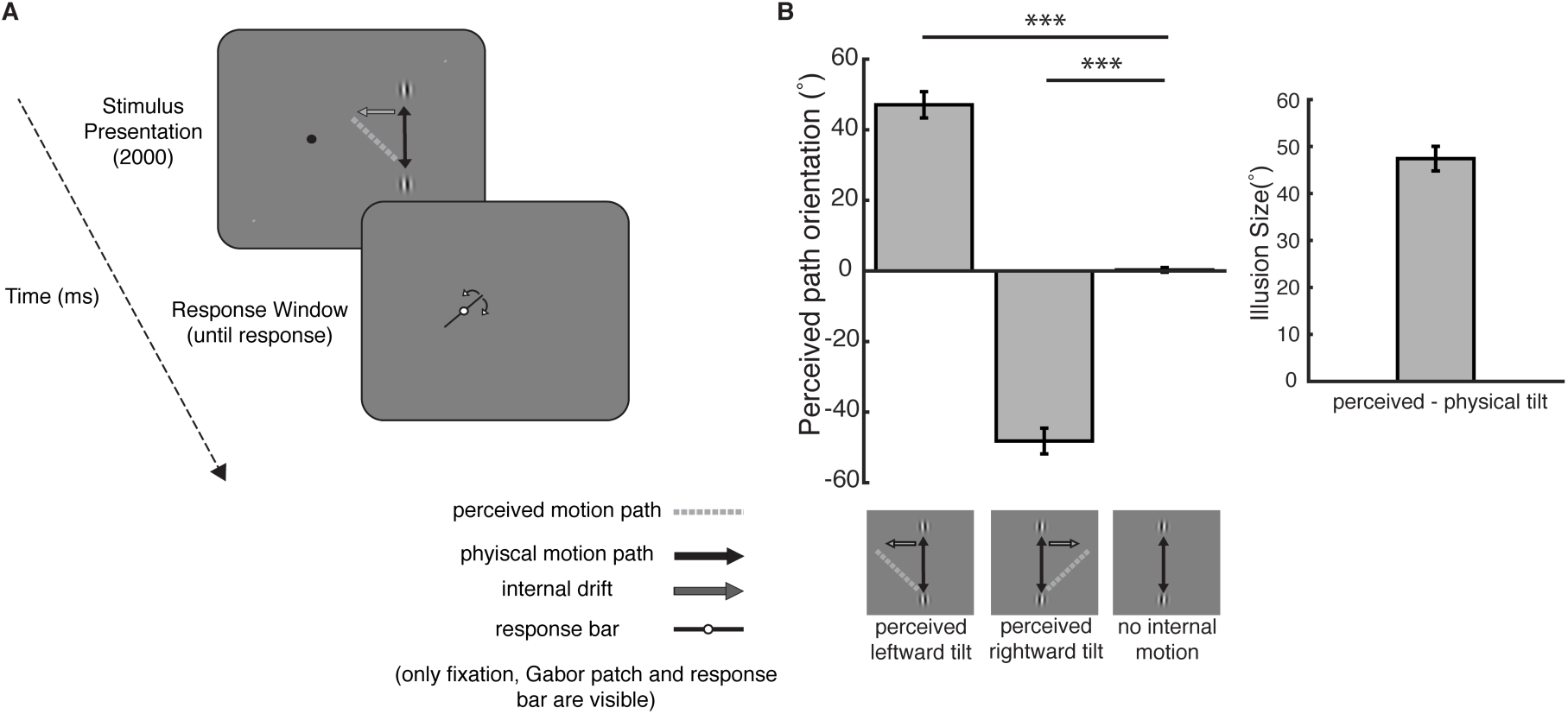
Behavioral task trial sequence and performance. (A) Each trial began with a Gabor patch shown in the right hemifield moving vertically (example stimulus is a double-drift stimulus with a possible perceived motion path tilted leftward driven by its internal motion) for 2 seconds which then disappeared. A response bar then appeared at fixation and remained on the screen until participants adjusted its orientation to the perceived motion path of the Gabor patch. (B) Group averaged perceived path orientation (°) of the double drift stimulus and control stimulus (no internal drift) and the illusion size of the double-drift effect. Error bars represent 95% CI.

### Perceived paths are decodable in V2 and V3 but do not share the same activation patterns with that of matched physical paths

We then used fMRI and MVPA to classify the activation patterns driven by a double-drift stimulus that moved along the same physical path but could produce two illusory paths with opposite perceived orientations depending on the direction of its internal drift motion. Importantly, as the internal drift of the double-drift stimulus reverses its direction at the two endpoints of the motion path, both illusory trajectories have equal periods of leftward and rightward local motion across a complete back and forth cycle, so the only difference between the two conditions is their perceived motion direction. We compared these perceived motion paths with those of matched Gabor stimuli, lacking internal drift motion, that physically moved in the direction of the two illusory paths as measured in the behavioral task for each subject (see **STAR Methods** for details). The MVP classification analysis was first conducted in voxels that showed significantly greater BOLD responses to the motion path locations within each of the early visual areas defined in a separate retinotopic mapping session. **Figure 3A** shows these 4 motion path ROIs (MPROIs) for V1, V2, V3, and MT+ on a representative participant (see **Table S1** for ROI sizes). When training the linear SVM classifier and testing on the left-out dataset from the same stimulus conditions, classification accuracies for the two control stimuli (physical leftward vs. rightward path orientation) were significantly above chance in all 4 MPROIs (*p*s < .001; *p* values were adjusted using FDR method in this and all subsequent analyses; **Figure 3B;** See **Table S1** for statistical results). This suggests that the activation patterns for the two stimuli with physical path orientations that matched those of the perceived path of the double-drift stimulus can be reliably differentiated in these retinotopic visual areas. However, classification accuracies for the two double-drift stimuli (illusory leftward vs. rightward path orientation) were significantly above chance only in area V2 (*p* = .004) and V3 (*p* < .001) but not in V1 (*p* = .44) or MT+ (*p* = .12) (**Figure 3B;** See **Table S1** for statistical results). This indicates that activity in V2 and V3 differed for the two perceptual motion paths of the double-drift stimulus whereas activity in V1 were indistinguishable for these two perceptual conditions. This is consistent with a recent study that finds evidence for the involvement of retinotopic areas with anatomically separated quadrantic representation of visual space, such as area V2 and V3, in deriving the illusory motion of this stimulus [17]. In addition, our results show that the activity pattern in MT+ did not correspond to the two perceived motion paths for the double-drift stimulus. This is surprising since there is evidence that suggests that neurons in MT+ represent the perceived position offsets driven by motion signals [18-20]. However, since the physical position was also not encoded as strongly in MT+ as in other early visual regions, e.g. V1-V3, it is possible that the lack of perceived position information in MT+ for this stimulus was due to weak position signals in this region.

**Figure 3.**
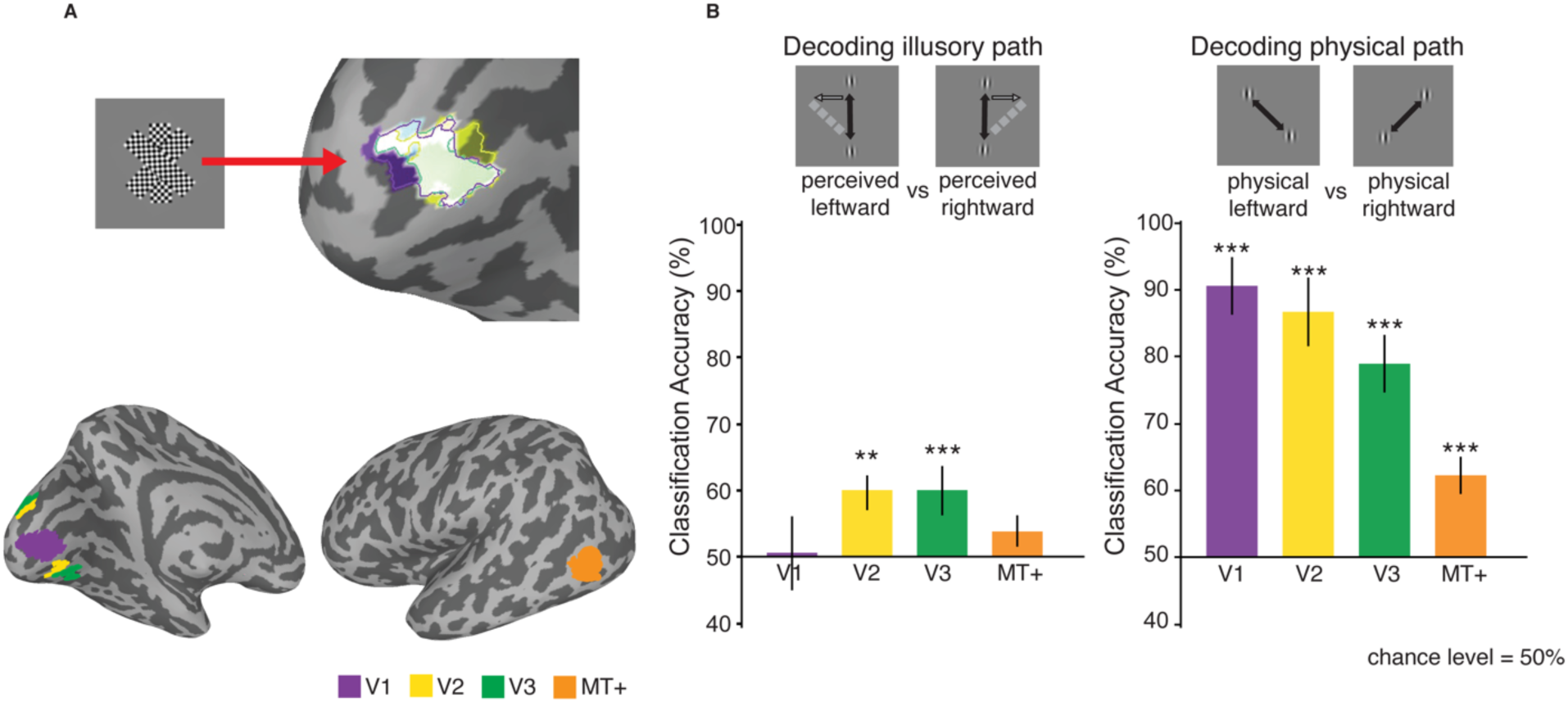
MPROIs and decoding accuracy. (A) Voxels for MPROIs were selected within each early visual area by combining regions that showed greater ctivation for any of the three tilted checkerboard rectangles than to fixation. Example ROIs shown in a presentative participant: V1 (purple), V2 (yellow), V3 (green), and MT+ (orange). (B) Classification accuracies on the two double drift stimuli (perceived rightward vs. leftward motion path) nd the two control stimuli (physical rightward vs. leftward moth path) in the MPROIs of V1, V2, V3 and T+. Error bars represent ± 1 SEM, ** *p* < .01, *** *p* <.001. See also Figure S3 and Table S1.

To directly examine whether the activation patterns for the illusory motion paths of the double-drift stimulus share a similar structure with those of the matched physical motion paths, we conducted a cross-decoding analysis where we trained the linear SVM classifier with the data corresponding to the two double-drift stimuli and tested with the data corresponding to the two control stimuli, as well as the reverse analysis where we trained the classifier on the control stimuli and tested on the double-drift stimuli. Interestingly, classification accuracies from cross-classification in either direction were not significantly different from chance in any of the MPROIs (*p* > .1; See **Table S1** for statistical results), including V2 and V3. Thus, although the activation patterns of the two double-drift stimuli can be differentiated in V2 and V3, their representations carried different information from those of their matched physical motion paths in these two areas.

### Representational structure in early visual areas reveals strongest dissimilarity between physically different motion paths

Since the previous analysis suggested different representations of illusory and physical motion paths in these early visual cortices, we further conducted representational similarity analysis (RSA) to examine the representational structure of the five stimulus conditions in the early visual MPROIs [21]. **Figure 4** shows the dissimilarity matrices (DSMs) of the five stimulus conditions V1, V2, V3 and MT+. Early visual areas V1-V3 exhibited the strongest dissimilarity between the stimuli with different physical motion paths (physical left path vs. rightward vs. vertical motion path) as compared to those that shared the same physical motion direction but with a large perceptual difference (double-drift stimuli: illusory leftward vs. rightward) (V1: r = 0.91, *p* = .001; V2: r = 0.81, *p* = .016; V3: r = 0.70, *p* = .05). This similarity structure confirmed that the representation of the double-drift stimulus in these early visual areas was largely influenced by its physical motion path. The representational structure in MT+ showed high similarity between all stimulus conditions (r = 0.27, *p* = .5).

**Figure 4.**
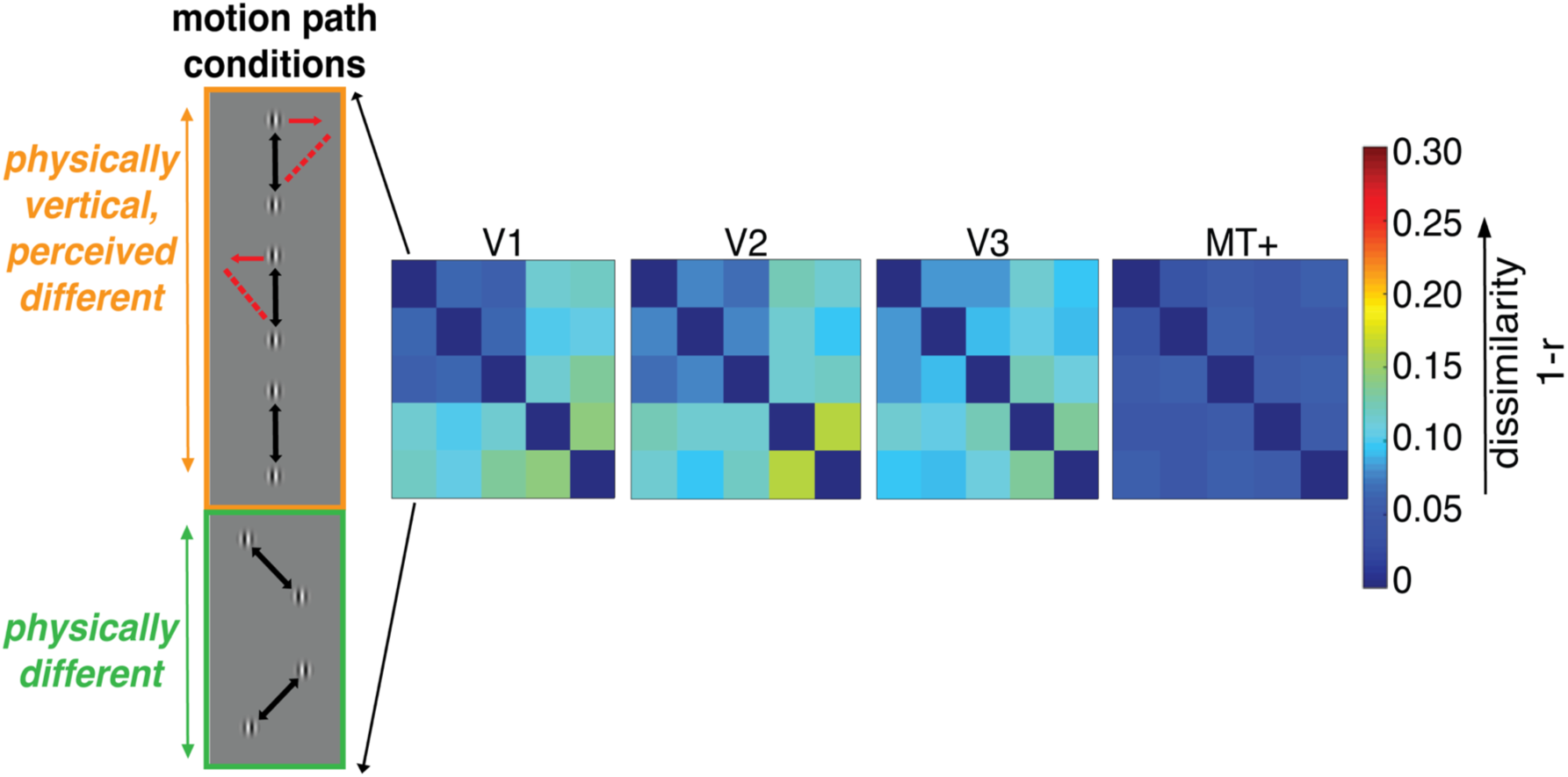
Results of the representational similarity analysis (RSA). Representational dissimilarity matrices for the five stimulus conditions in MPROIs (V1, V2, V3 and MT+).

### No difference in BOLD response amplitude between illusory and matched physical paths

We also calculated BOLD signal changes of each stimulus condition relative to baseline within each MPROI to examine whether the above-chance decoding accuracies for the illusory and physical motion paths from MVPA could be detected at the univariate level. Group averaged BOLD time courses corresponding to the illusory and matched physical motion conditions from each MPROI are shown in **Figure S3**. All early visual MPROIs exhibited above-baseline activity for the five stimulus conditions (*p*s < .05) except V1, which showed above-baseline activation only for the control stimulus with leftward motion direction (*p* < .05). Importantly, we observed no difference in response magnitude for the two double-drift stimuli (*p*s > .1), or for the two control stimuli with matched physically different motion paths in these MPROIs (*p*s > .1), suggesting that the above-chance decoding accuracies in these regions from MVPA cannot be simply explained by differences at the aggregate activation level. In addition, there was no significant difference in mean signal intensity between the double-drift and the control stimulus that had the same physical path direction but differed in the presence of internal motion in any of these MPROIs (**Figure S3C**; *p*s > .1), suggesting that the two conditions were matched in terms of stimulus energy. Thus, the failure in cross-decoding in these regions was not simply due to a mismatch of internal motion in the double-drift and control stimulus.

### Higher-order regions show a shared representation of the illusory and matched physical motion paths

Our classification results showed that the illusory motion path can be reliably decoded in some of the early visual MPROIs. To further explore areas in the brain that could decode the illusory motion paths beyond the pre-defined visual ROIs, we conducted a whole-brain searchlight analysis using a 4-voxel radius spherical searchlight. The same decoding analyses for the illusory and matched physical paths as for the ROI-based analysis were conducted, and results of the searchlight analysis were corrected using a cluster thresholding method for multiple testing (see **STAR Methods** for details). **Figure 5** shows the group accuracy maps from the classification searchlight analysis for decoding the illusory motion paths (**Figure 5A**) and the matched physical paths (**Figure 5B**). We identified several clusters in the two hemispheres that showed decoding accuracies that were significantly higher than chance levels for the illusory motion path outside our pre-defined early visual ROIs: a large cluster that spans the superior frontal and medial frontal gyrus, and several clusters in the superior temporal gyrus, dorsal anterior cingulate cortex, and the left postcentral gyrus. We also found a cluster spanning the early visual cortex that confirmed our significant decoding results in the ROI-based analysis (see **Table S2** for a complete list of significant clusters). Decoding the matched physical motion paths yielded an even larger range of cortical regions, including visual, parietal, and frontal areas (**Table S2**).

**Figure 5.**
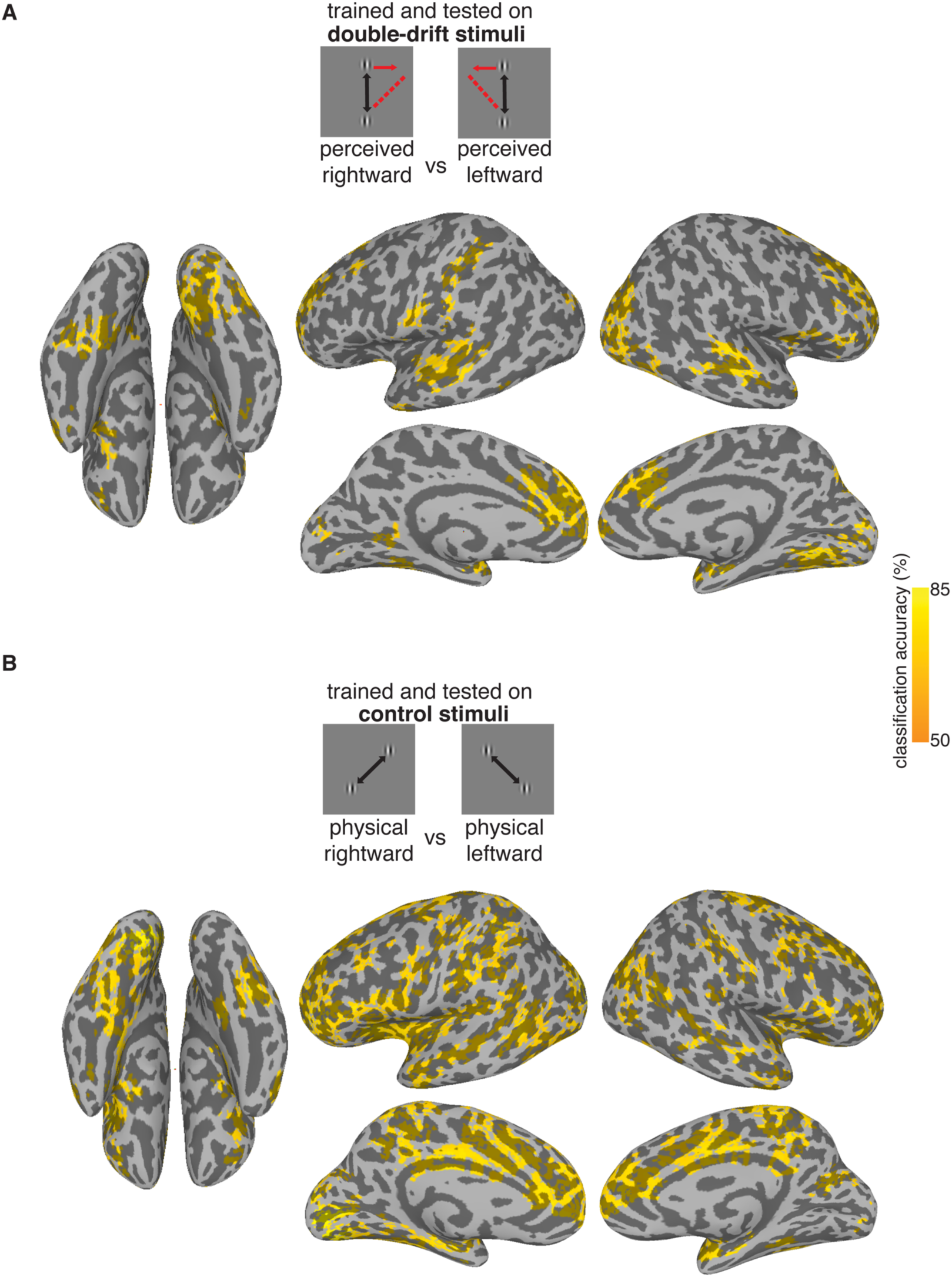
Group accuracy maps of the within-condition classification searchlight analysis. (A) Decoding the illusory motion paths. (B) Decoding the matched physical motion paths. Results were thresholded at *p* = .01 and FDR corrected across clusters at *p* < .05. See also Table S2.

Our ROI-based cross-decoding results showed that the activation pattern for the double-drift stimulus had little or no similarity to that of its matched physical motion path in any of the localized regions of the early visual areas. It is however possible that a shared encoding of the illusory and matched physical motion path of the double-drift stimulus is represented somewhere outside our pre-defined visual ROIs. This shared encoding could be a marker of the emergence of the perceptual, as opposed to the retinal, location of the double-drift stimulus. We therefore also conducted a whole-brain searchlight analysis using a cross-decoding classifier between the double-drift and control conditions to further explore the locus of any such shared representations. The results of this searchlight analysis should yield regions with similar patterns of activation for the double-drift and control stimulus that have the same perceived orientation. Interestingly, we found several significant clusters in anterior parts of the brain that have above-chance cross-decoding between the illusory and matched physical paths (**Figure 6;** see **Table S3** for a complete list of significant clusters) but, in agreement with the previous ROI analysis, none in early visual areas. Specifically, we found clusters that had above-chance cross-decoding in both directions (i.e. trained on double-drift then tested on control stimuli and trained on control then tested on double-drift stimuli) in the anterior cingulate and medial frontal gyrus in both hemispheres, anterior part of the middle frontal gyrus in the left hemisphere, left inferior parietal lobule and parahippocampal gyri. Besides these overlapping regions, cross-decoding from double-drift to control stimuli resulted in additional significant clusters in the middle and inferior frontal gyrus and medial frontal gyrus in both hemispheres; cross-decoding from control to double-drift stimuli produced additional significant clusters in the right precentral gyrus and left parahippocampal gyrus. In addition to these cortical clusters, we also found several subcortical clusters as detailed in **Table S3**.

**Figure 6.**
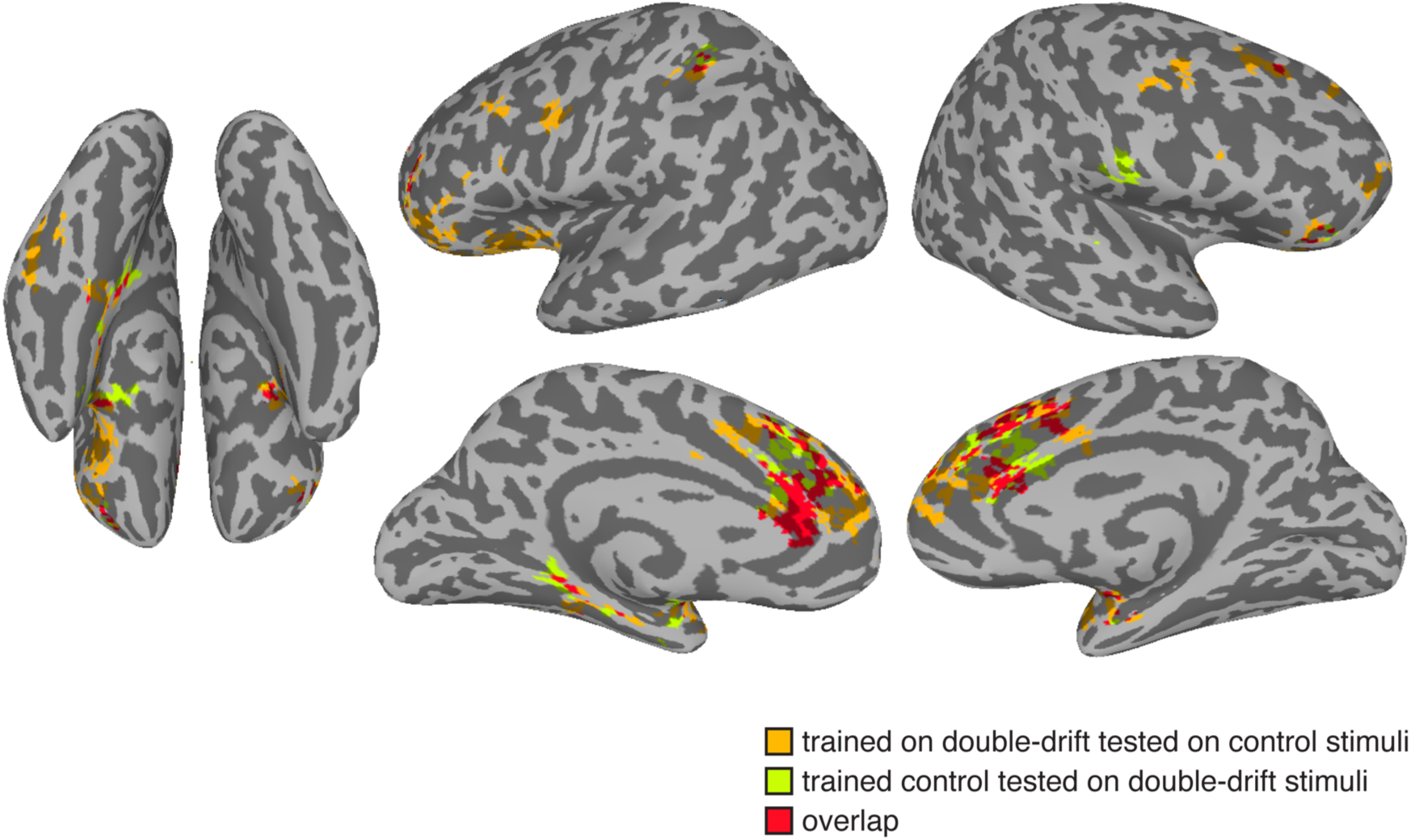
Cluster-thresholded searchlight map with significant above-chance cross-decoding accuracy. Orange represents significant clusters when training the classifier on double-drift and tested on the control stimuli. Green represents significant clusters when training the classifier on the control and tested on the double-drift stimuli. Results were thresholded at *p* = .01 and FDR corrected across clusters at *p* < .05. See also Figure S4 and Table S3 and Table S4.

To exclude the possibility that the failure in cross-decoding in regions such as the early visual cortex was caused by (potentially) subtle difference in mean signal intensity across conditions, even though their differences in mean activation amplitude were not statistically significant (**Figure S3**), we performed an additional searchlight analysis where we removed the grand mean of each stimulus condition within each searchlight. The results remained qualitatively similar to that of the original cross-decoding searchlight analysis as shown in **Figure 6**, with no significant clusters observed in early visual cortex (see **Figure S4** and **Table S4**). This suggests that failure in cross-decoding in regions such as the early visual cortex was not simply due to a difference in stimulus-driven responses between stimulus conditions of the training (e.g. physical leftward vs. rightward paths) and testing dataset (e.g. illusory leftward vs. rightward paths).

## Discussion

The aim of this experiment was to localize areas of cortex associated with perceived object positions and trajectories when they differ from the positions and trajectories registered on the retina. Normally, the location of retinal input and the corresponding perceived positions are well matched, but in a remarkable motion-induced position shift, the ‘double drift’ illusion, there is a dramatic mismatch between physical and perceived position. In particular, the perceptual displacement can be as large as several degrees of visual angle and can build for a second or more [1].

Although we found that the perceived motion path of the double-drift stimulus can be decoded in early visual areas V2 and V3, results from cross-classification analysis reveal that the activation patterns that differentiated the illusory motion paths in these early visual regions were not related to the activity patterns that encoded the physical motion paths with matched perceptual orientation. Therefore, the basis of our classification results in these early regions is likely not related to the perceived motion path *per se* but might arise from lower level properties of the stimulus such as the combined vector of the local and global motion. For example, leftward internal drift was associated with an upward external motion in one case (e.g. double-drift stimulus with perceived leftward trajectory), and with a downward external motion in the other (e.g. double-drift stimulus with perceived rightward trajectory).

Interestingly, the significant cross-classification clusters found in our searchlight analysis were primarily in areas that are known to be involved in executive control, such as the lateral prefrontal cortex (LPFC) [22,23], dorsal anterior cingulate (dACC) (the cingulo-opercular control network; [24,25]), pre-supplementary motor area (preSMA) [26] and medial prefrontal cortex (MPFC) [27], and in spatial information processing such as the inferior parietal lobule (IPL) [28,29]. This indicates that the neural coding for the perceived motion path in these high-order regions was driven by a representation of the illusory motion path that was similar enough to that of the matched physical motion path to permit cross-decoding. Importantly, these regions have been implicated in the literature in transforming sensory representations to different representational formats for different functional purposes. For example, lateral frontal regions have been implicated in forming abstractions of incoming information [30,31]. Motor-related areas such as preSMA are involved in encoding past and current information for perceptual decision making and generating motor preparatory signals from readout of sensory information [32-34]. The cingulo-opercular network was shown to be involved in downstream control process for perceptual recognition and working memory output gating, by integrating information accumulated from the frontoparietal and sensory regions [35,36]. IPL has been implicated in transforming visuospatial information into motor output [37-40]. Therefore, successful cross-classification in higher-order cortical regions as observed in our study may reflect similar representational changes from a sensory format to a different, more abstract format, which could allow for generalization between different physical stimuli in a shared format of perceptual experience.

In comparison, representations in early visual cortex are sensitive to stimulus-specific changes and therefore may not permit successful cross-classification between the illusory and matched physical motion paths. Indeed, our results show that this high-level representation of perceived as opposed to real stimulus positions was not shared with or projected back down to early visual areas. Instead, the observed above-chance classification in V2 and V3 for the perceived motion path might suggest that activity in these areas encodes the combined motion signals integrated over short durations. These local direction errors are the base data that get integrated into the illusory path, but likely do not account for the illusion alone, because these errors appear to accumulate over long durations, and cells in these early processing stages do not have the second-long integration time windows that the double-drift stimulus requires to build up the position deviations. Indeed, one distinct feature that makes this illusion such a powerful effect is that its perceived motion path can be formed by accumulating position shifts over long durations of a second or more, while other motion-induced position shifts effects like the flash-grab stimulus, only integrate motion signals over about 90 ms [10]. Given the short decay time constant for orientation cells in early visual areas [41], it is possible that a motion-position integration process of such a long duration requires higher-order brain areas to store and accumulate position offsets in order to form a consistent motion trajectory. Thus, our results suggest that the higher-order areas where we find significant cross-classification could be candidate areas that accumulate outputs from V2 and V3 so that perception continues to drift farther away from the real path for over a second.

Our finding that there is no shared activity in early visual cortex that corresponds to the physical and perceived paths conflicts with some of the previous fMRI studies of motion-induced position shifts [19,42,43]. In the case of the “flash-grab” illusion where a flash is pulled forward by the motion that underlies it [10], it was shown that neural activity for the perceived position shifts of this stimulus correlates strongly with activity seen for physical stimuli with locations matched to the illusory ones solely in early visual cortices V1 through V3 but not in higher-order areas [43]. Similarly, for the related “flash-drag” illusion [44], activity in early retinotopic cortical areas, most notably MT+, also shows strong correlation between perceived position of a flash that was shifted by surrounding moving patterns and matched physical positions [19]. We suggest that the involvement of top-down attentional signals may account for the discrepancies between these results and ours. In particular, when higher-order areas generate a percept of an object, downward attentional signals can feed back to early visual cortex and generate activation at locations where the object is expected, rather than where it is in retinotopic coordinates [45]. These downward signals would complicate any attempt to determine where perceptual representation begins as activations in early cortical areas would be composed of a combination of bottom-up activations matching the physical stimulus, and top-down activations that, based on the percept, matched its expected location and properties. The presence of such top-down activations would support cross-classification between perceived and matched physical motion paths in all areas that received such feedback. Although there is not yet direct evidence that attention is extrapolated to match these shifts in perceived position, it is well established that saccades are directed to the perceived rather than physical locations for the motion-induced position shifts that displace the target along the direction of motion, such as the flash-lag and flash-drag stimuli [46-49]. Given the close link between the saccade and attention system [50,51], it is reasonable to assume that attention will be shifted by these illusions to the same extent as perception and saccades.

For the double-drift illusion, on the other hand, saccades are directed to the physical location of the stimulus, not its perceived location [1]. The dissociation of saccades and perception is unique to this stimulus as other motion-induced position shifts affect saccades and perception equally [46-49]. Given the immunity of saccades to the double-drift illusion, we speculate that attentional shifts, like saccades, are not affected by the illusion either. If correct, any downward projections from areas involved in attentional shifting circuitry would prioritize the stimulus’s physical locations, rather than its perceived, illusory ones. If this is the case, there would be no activity at cortical regions corresponding to illusory locations prior to those areas in the visual processing hierarchy that actually do encode the perceived location. Since we find no cross-classification between perceived and matched physical paths in early visual areas for the double-drift stimulus, we assume that either such attentional feedback is weak if it is to the perceived locations, or more likely, that the top-down feedback is in fact to the physical locations. If this assumption is correct, this stimulus affords the unique possibility of probing where the perceptual coordinates of object position arise in the processing hierarchy without the confound of top-down attentional projections. The feedback signals in this case would simply prioritize the stimulus’s representation in physical instead of perceived coordinates and so would not mask the point at which perception deviates from the bottom up signals.

In summary, our data place a lower limit on where areas of the brain are located that are in perceived as opposed to retinotopic coordinates. Remarkably, there was no cross-classification of corresponding real and perceived motion paths in early visual areas, a finding probably linked to the absence of influence of the illusion on saccades, and by inference on attentional shifts as well. Without downward projections activating the perceived locations, cortical regions prior to the emergence of the percept then show only the uncontaminated bottom-up activity. And this reveals that the representation of perceived position likely emerges much later in the processing hierarchy than in early visual cortical areas, even if early areas such as V2 and V3 provide the instantaneous direction errors that are then integrated in later areas to compute the final percept. To the extent that visual consciousness coincides with perceived position, our data also place clear constraints on the neural correlates of visual consciousness, at least for this particular type of stimulus.

## Acknowledgments

The research leading to these results received funding from the Department of Psychological and Brain Sciences (PC and PT), NSF grant 1632738 (PT), a Templeton Foundation Grant 14316 (PT) and an NSERC Canada grant (PC).

## Author contributions

S.L., Q.Y., P.U.T. and P.C. designed the experiment. S.L. and Q.Y. conducted the experiment and analyzed the data. Q.Y. contributed unpublished analytic tools. P.U.T. and P.C. supervised the entire project. S.L., Q.Y., P.U.T. and P.C. wrote the manuscript.

## STAR Methods

### 1. CONTACT FOR REAGENT AND RESOURCE SHARING

Further information and requests for resources and reagents should be directed to and will be fulfilled by the Lead Contact, Sirui Liu.

### 2. EXPERIMENTAL MODEL AND SUBJECT DETAILS

#### 2.1 Subjects

Nine individuals from the Dartmouth College community participated in this study (5 females; age range: 21-32, mean age = 26.6 +-3.1). All participants were naïve to the purpose of this study and had normal or corrected-to-normal vision. Written, informed consent approved by the Committee for the Protection of Human Subjects at Dartmouth College was obtained from each participant prior to the first experimental session. Participants were screened by the Dartmouth Brain Imagining Center fMRI Subject Safety Screening Sheet and received a compensation of $20/hour.

### 3. METHOD DETAILS

#### 3.1 Stimuli

All stimuli were generated using Matlab 2015a [52] and PsychToolbox-3 [53]. The stimulus in the behavioral and the main fMRI experiment was based on the double-drift stimulus used in [1] (see **Figure 1** and **Movie S1**). We used a Gabor pattern (sinusoidal grating within a Gaussian envelope) with a spatial frequency of 0.5 cycle/dva (cycles per degree of visual angle) and 100% contrast presented on a uniform gray background (53 cd/m^2^). The standard deviation of the contrast envelope was 0.4 degree of visual angle [dva]. The Gabor pattern moved back and forth along a linear path of length 5 dva, with a speed of 5 dva/sec (external motion). The sinusoidal grating had the same orientation as the motion path, and drifted in the orthogonal direction with a temporal frequency of 4 Hz (internal motion), reversing direction at the two endpoints every 1 second (‘double-drift stimulus’), or stayed static during the trial (‘control stimulus’). The midpoint of the trajectory was placed at 5 dva to the right of the screen center. A black fixation point (0.3 dva diameter) was presented at 3 dva horizontal to the left of the screen center throughout all the behavioral and MRI experiments. We moved the fixation to this location so that our stimulus was 8 dva away from fixation. This was the eccentricity at which [1] found a large perceptual effect. In the pre-scan behavioral task, participants reported the perceived orientation of the motion path using a black line (‘response bar’) centered at fixation that was 0.05 dva in width and 5 dva in length.

#### 3.2 Pre-scan behavioral task

Stimuli were presented on an Apple iMac Intel Core i5 (Cupertino, CA) and were displayed in a dark room on a 16’’ ViewSonic G73f CRT monitor (1024 × 768 pixels at 90-Hz) placed 57-cm from the participant with their head stabilized on a chinrest during the experiment. **Figure 2A** shows a sample trial of the pre-scan behavioral task. Participants were instructed to keep their gaze at the fixation point throughout the experiment. In each trial, a Gabor patch was shown in the periphery and moved back and forth in a vertical path for 2 s. Its internal texture drifted either leftward, rightward or remained static. For each participant, the drift direction of the internal texture was randomized across trials. Following the stimulus, participants were instructed to rotate the response bar by pressing the corresponding keyboard keys (up arrow for counterclockwise; down arrow for clockwise) until its direction matched the perceived angle of the motion trajectory of the Gabor. The response bar was presented at a random orientation for each trial and remained on the screen until participants were satisfied with their response and pressed the space bar for the next trial. Overall, each participant completed ten adjustment trials for each internal drift direction for a total of 30 trials that lasted about 15 minutes.

#### 3.3 MRI acquisitions

The scanning was conducted on a 3T MRI scanner (Philips Intera Achieva) with a 32-channel head coil at the Dartmouth Brain Imaging Center at Dartmouth College. For each subject, we collected functional BOLD activity using an echo-planar imaging (EPI) sequence (TR = 2 s, TE = 35 ms, voxel size = 3 × 3 × 3 mm, flip angle = 90°, FOV = 240 × 240 mm) and a high resolution anatomical scan using T1-weighted MPRAGE sequence at the end of each scanning session (voxel size = 1 × 1 × 1 mm).

#### 3.4 Main experiment runs

Stimuli were presented on a screen (47.5 cm width) at the back of the scanner through an LCD projector. The screen resolution was 1024 × 768 pixels with 60 Hz refresh rate. The projected stimuli were viewed through a mirror located on the head coil with a viewing distance of 101.6 cm. Participants completed 10 fMRI main experimental runs. In each run, after an initial 4 s blank fixation period, participants viewed a total of fifteen trials, each of which was composed of a 11s stimulus block followed by a 15s fixation block with the order of the trials randomized for each participant. Each stimulus block was composed of a Gabor patch presented in the right hemifield that moved back and forth along a linear path for five repetitions (2s each repetition) and then disappeared for 250 ms in between repetitions. In total, each experimental run was 394s long. **Figure S1A** illustrates the five stimulus conditions used in the main fMRI experiment. Participants viewed three blocks per stimulus condition in each run. To make sure participants were attending to the stimulus, the contrast of the presented Gabor stimulus reduced 50% randomly in each run for 200 ms and participants were asked to press a response button each time they saw the change. **Figure S1B** shows a sample trial sequence for the main fMRI main experiment. We also conducted two additional EPI runs using a rectangular checkerboard pattern flickering at 8 Hz that covered the spatial extent of the perceived and physical motion paths of the double-drift stimulus. The checkerboard pattern was centered at 8 dva horizontal to the right of the fixation with its height the length of the motion path of the Gabor pattern and a width the size of the Gabor stimulus. **Figure S2A** shows the three conditions in the stimulus location localizer runs: vertical, leftward or rightward tilted checkerboard rectangle. The two oblique checkerboard stimuli were tilted in the direction of the perceived motion path for the double-drift stimulus. The tilt angle was individually calculated from the responses in the pre-scan behavioral task for each subject. Each run contained an initial 4s fixation block and fifteen trials, each of which was composed of a 10s-stimulus block with a flickering checkerboard pattern followed by a 12s. blank fixation block. There were five trials per stimulus condition for a total of fifteen trials per run with the order of the blocks randomized for each participant. **Figure S2B** shows the trial protocol. To maintain attention to the stimulus, participants were asked to press a response button each time they saw the color of the fixation point changed.

#### 3.5 Region-of-interest localization runs

In addition to the main experiment, participants completed a separate scanning session that included a standard retinotopic mapping procedure and three MT+ localizer runs (292 s each). We followed the standard retinotopic mapping procedure [54,55] by using clockwise or counterclockwise rotating checkerboard wedges (flickering at 4 Hz, ten 192 s runs) to map polar angle and using expanding or contracting checkerboard rings (flickering at 4 Hz, four 192s runs) to map eccentricity. The fixation point in our experiment was moved 3 dva horizontal to the left of the center of the screen to match that of our main experiment. MT+ was functionally localized for each participant following the procedure from [56]. In each run, participants viewed seven 16s blank fixation blocks interleaved with six 30s stimulus blocks. The stimulus was composed of one hundred 0.3dva diameter black dots spanning the whole visual field that either moved coherently, flickering at 30 Hz, or remain static on the screen. Each of the three stimulus conditions was presented twice in each run with the order randomized for each subject. For the coherently moving condition, the dots could be moving rightward, leftward, vertically upward or downward, expanding, contracting, rotating clockwise or counterclockwise at 7 dva/s with 100% coherence, while resetting their locations every 367 ms. To make sure they were fixating, participants were asked to press a response button each time they saw the color of the fixation point changed for all of the localizer runs.

### 4. QUANTIFICATION AND STATISTICAL ANALYSIS

#### 4.1 Behavioral data analysis

For each participant, we first determined the perceived path angle of the double-drift stimulus (with leftward or rightward perceived motion paths) and the control stimulus that lacked internal motion away from the physical, vertical orientation. Paired-samples t-tests were conducted using R and RStudio v.1.0.136 to compare the mean differences of the perceived path orientation between the double-drift stimulus and the control stimulus with no internal drift [57,58]. The magnitude of the double-drift illusion was then calculated individually by taking the difference between these two measurements. A positive value of the illusion size indicates that the perceived motion orientation was biased toward that of the internal drift. The average of this value was then used in the following scanning session as the motion direction for the control stimuli that moved obliquely with no internal drift as well as the tilt angle for the checkerboard pattern in the localizer runs to define ROIs for the motion path of the stimulus for each subject.

#### 4.2 fMRI data analysis

##### 4.2.1 Preprocessing

Functional imaging data was preprocessed using AFNI [59]. For each participant, the EPIs were first registered to the last run of each scan session and then motion corrected, linearly detrended, and z-scored within each run. The anatomical images collected in the first scanning session were aligned to the EPI scans of the same session. Anatomical scans collected in the two localizer runs that define motion path locations in the second scanning session were first aligned to that of the first scanning session before aligning to the EPI scans. Localizer data were further smoothed with a 4-mm FWHM Gaussian kernel. For the searchlight analysis, the EPI scans were normalized to the Talairach standard space [60].

##### 4.2.2 ROI definition

The cortical surface of each participant was first reconstructed with FreeSurfer [61] using the high-resolution anatomical images. All data in the localizer runs were first mapped onto this cortical surface to define the ROIs. Early visual areas left V1, V2, and V3 were individually drawn by hand on individual surfaces based on the phase angle maps computed from data in the retinotopic mapping session. MT+ was individually defined on surface based on data from the MT+ localizer runs using beta coefficient values calculated from a General Linear Model (GLM) analysis that specified voxels that responded more strongly to moving than to stationary dot patterns (*p* < 10^-4^ after correcting for multiple tests using false-discovery rate (FDR) [62]. To identify the voxels that responded to the motion path of the double-drift and control stimulus within each of the ROIs, we then selected the voxels that showed significantly greater activation for any of the three tilted checkerboard rectangles than to fixation (*p* < 10^-4^, FDR corrected) in the left hemisphere and only these voxels were included for the rest of the ROI-based analysis. All these surface-defined results were then individually mapped back into the volume space and aligned to the EPI data of the first scanning session by aligning the anatomical scans of the two sessions for subsequent analysis.

##### 4.2.3 Time course of BOLD activity

To create the time series of BOLD activity in each ROI, we averaged BOLD activity in all voxels within the ROI and calculated the percent signal change in activation relative to baseline for each TR of each trial. Baseline was defined as the activation of the first TR of each trial. Average BOLD activity at each time point was then calculated by averaging the percent signal change across trials within each condition. One-sample t-tests against 0 were used to assess statistical significance above baseline for each TR within each ROI for each stimulus condition at *p* < 0.05 after correcting for multiple tests using FDR [62]. In addition, paired samples *t*-tests were used to compare BOLD activity between 1) the two double-drift stimulus conditions, 2) the two control stimulus conditions, and 3) the double-drift stimulus conditions with internal motion and the vertically moving control stimulus without internal motion at each time point within each ROI and significance of the tests were determined at *p* < 0.05 after correcting for multiple tests using FDR [62].

##### 4.2.4 Multivariate pattern analysis (MVPA)

All the subsequent analyses were performed using the PyMVPA toolbox [63]. We first used PyMVPA to perform MVPA within each ROIs. For each trial, we extracted raw data averaged for 6 to 14 s after trial onset (considering a 6-s of hemodynamic delay) and fed the averaged data into linear support vector machines (SVMs) to implement classification of stimulus conditions. We performed two types of classification analyses: the first analysis was to classify between the two physically different motion paths of the control stimulus, or between the two illusorily-different motion paths of the double-drift stimulus, using a leave-one-run-out cross-validation procedure. To examine whether the activation patterns of the double-drift stimuli resembled that of the corresponding control stimuli, a second cross-decoding analysis was conducted using the same data, except that the training and test data were from separate conditions (i.e., training with the data corresponding to the control stimuli with matched physical motion path and testing with the data corresponding to the double-drift stimuli with physically vertical but perceived different motion path; and vice versa). Statistical significance of classification accuracies across subjects was determined by randomly shuffling the stimulus condition labels 1000 times to construct null distributions for each ROI and testing for significance above chance at *p* < .05 after correcting for multiple tests using false-discovery rate (FDR) [62].

The SVMs were further combined with a spherical searchlight procedure for whole-brain classification analysis. Specifically, we applied a volume-based searchlight analysis by sliding a 4-voxel-radius spherical linear SVM classifier voxel-by-voxel over the whole brain. As with the ROI-based analysis, the searchlight analysis was performed for decoding illusory paths, the matched physical paths, and cross-decoding using a leave-one-run-out cross-validation procedure. Group-level statistical significance for the searchlight analyses was determined following a cluster thresholding approach [64]: 100 permuted searchlight accuracy maps were first generated for each subject by randomly permuting the stimulus condition labels across trials. Then 100,000 group-average accuracy maps were computed by randomly sampling from each subject’s permutated maps to construct a null distribution of accuracy values. These bootstrapped average maps were then thresholded at *p* = 0.01 per voxel and were used for cluster-forming and for constructing the null distribution of cluster sizes for testing the significance of the real group-average map’s clusters. Significance of the test was determined at *p* < 0.05 across clusters of size larger than 30 voxels after correcting for multiple comparisons using the FDR [62]. The same set of searchlight analyses were performed where the grand mean of each stimulus condition was removed within each searchlight following the same cluster-based permutation tests and multiple comparison correction methods described above. Results were projected to the cortical surface reconstructed from the Talairach template [60].

##### 4.2.5 Representational similaritiy analysis (RSA)

To examine the neural representational geometry of the stimulus conditions, we also conducted a representational similarity analysis (RSA) [21]. This was done by calculating the Euclidean distance between patterns of responses for different stimulus conditions. For each ROI, a dissimilarity representational matrix (1-similarity) for the five stimulus conditions was derived. We then conducted a correlation test between each of these dissimilarity matrices with a hypothesized correlation pattern that corresponds to the case when the stimulus conditions with physically different motion path produce strongest dissimarlity (i.e. 1-similarity = 1) and those with physically same but perceptually different motion path (i.e. double-drift stimuli) were least dissimilar (i.e. 1-similarity = 0). P-values were then corrected for multiple tests using FDR [62].

### 5. DATA AND SOFTWARE AVAILABILITY

Custom analysis codes and fMRI data from the experiment are available upon request.

### Supplemental items

**Movie S1. Related to Figure 1**. Double-drift stimulus. This movie shows an example of the double-drift stimulus used in the experiment. The Gabor patch moves back and forth on a linear vertical path at the right hemifield. When fixating at the black dot, the internal grating drifts orthogonally toward the left as the Gabor patch moves upward and reverses its direction at the path reversal. The internal motion of the Gabor patch drives its perceived path to appear tilted leftward rather than vertical.

## Supplementary Information

**Figure S1.**
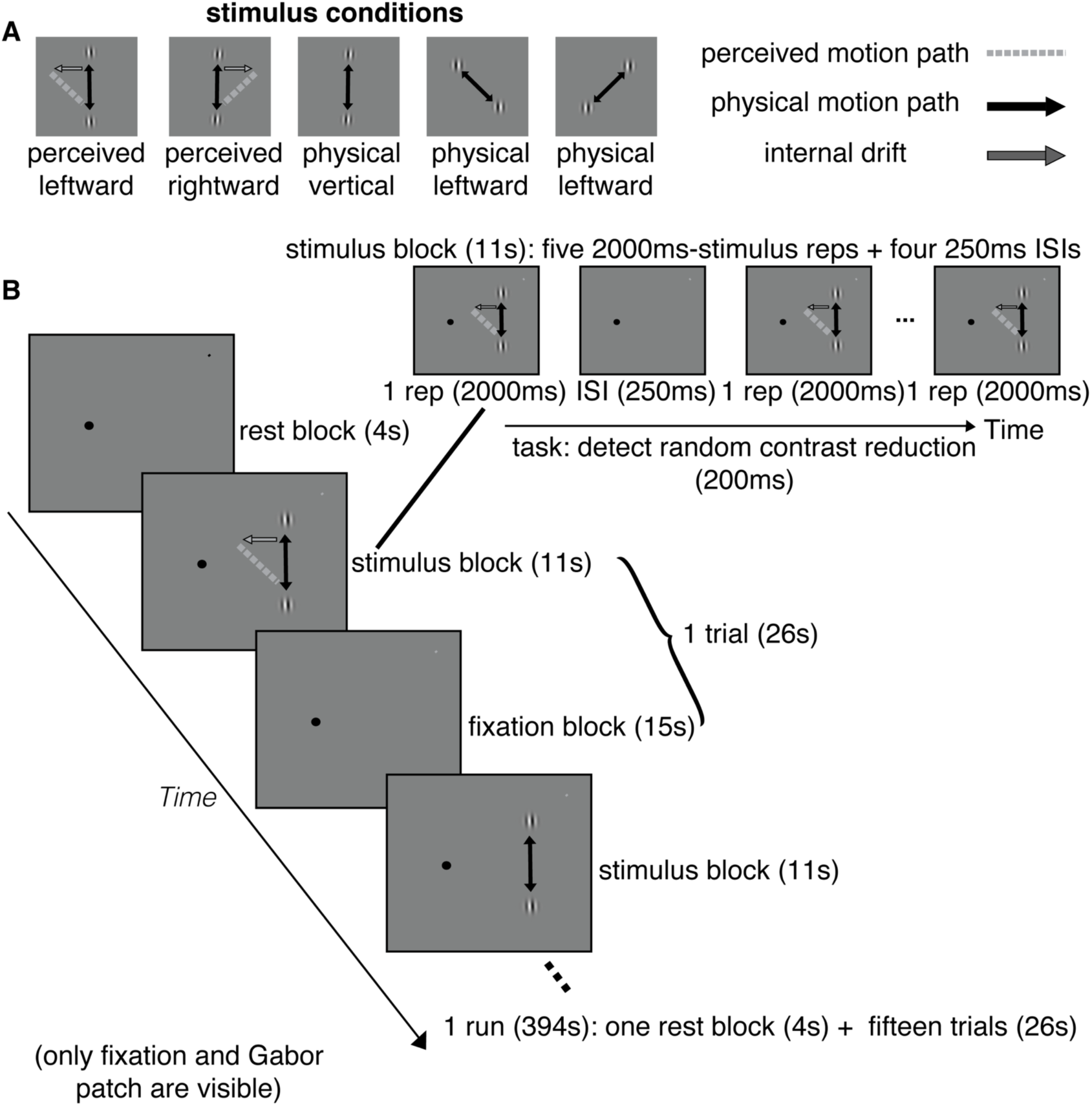
Related to STAR Method. Stimulus conditions and study protocol of the main fMRI experiment. (A) Stimulus conditions. The double-drift stimuli had a vertical physical motion path with opposite internal drift directions that could make the perceived motion path appear rotated either leftward or rightward relative to the physical motion path. The three control stimuli had either vertical, leftward or rightward physical motion paths with no internal drift. (B) Main fMRI experiment protocol: Each run lasted 394s and started with a 4s fixation block followed by fifteen repetitions of 26s stimulus trials. Each trial was composed of an 11s stimulus block followed by a 15s fixation block. In each stimulus block, participants viewed a moving Gabor patch in the right hemifield (example shows a double-drift stimulus with leftward perceived motion path due to the internal drift) presented for 2s for a total of five repetitions with a 250 ms ISI in between repetitions. Participants were asked to detect a brief contrast reduction (200 ms) of the Gabor patch presented at a random moment during each trial.

**Figure S2.**
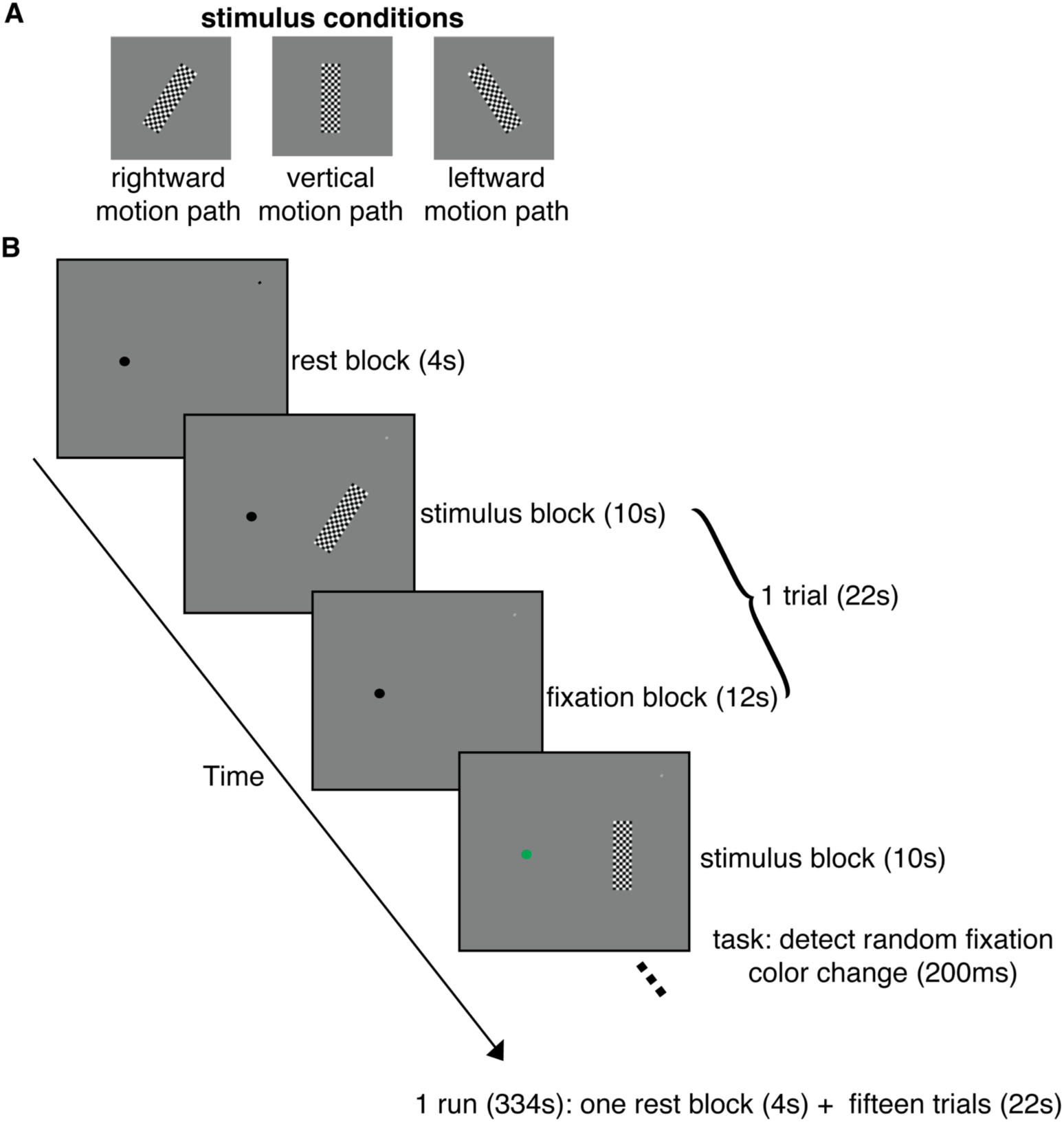
Related to STAR Method. Stimulus conditions and study protocol of the fMRI localizer experiment. (A) Stimulus conditions. The checkboard pattern had three orientations that matched the physical (i.e. vertical motion path) or the measured perceived path orientation of the double-drift stimulus (i.e. rightward/leftward motion path). (B) Each localizer run lasted 334s and started with a 4s rest block followed by fifteen stimulus trials (22s). Each trial was composed of a 10s-stimulus block with the stimulus flickered at 8 Hz followed by a 12s fixation block. Subjects were asked to detect random color changes (200 ms) of the fixation during the scan.

**Figure S3.**
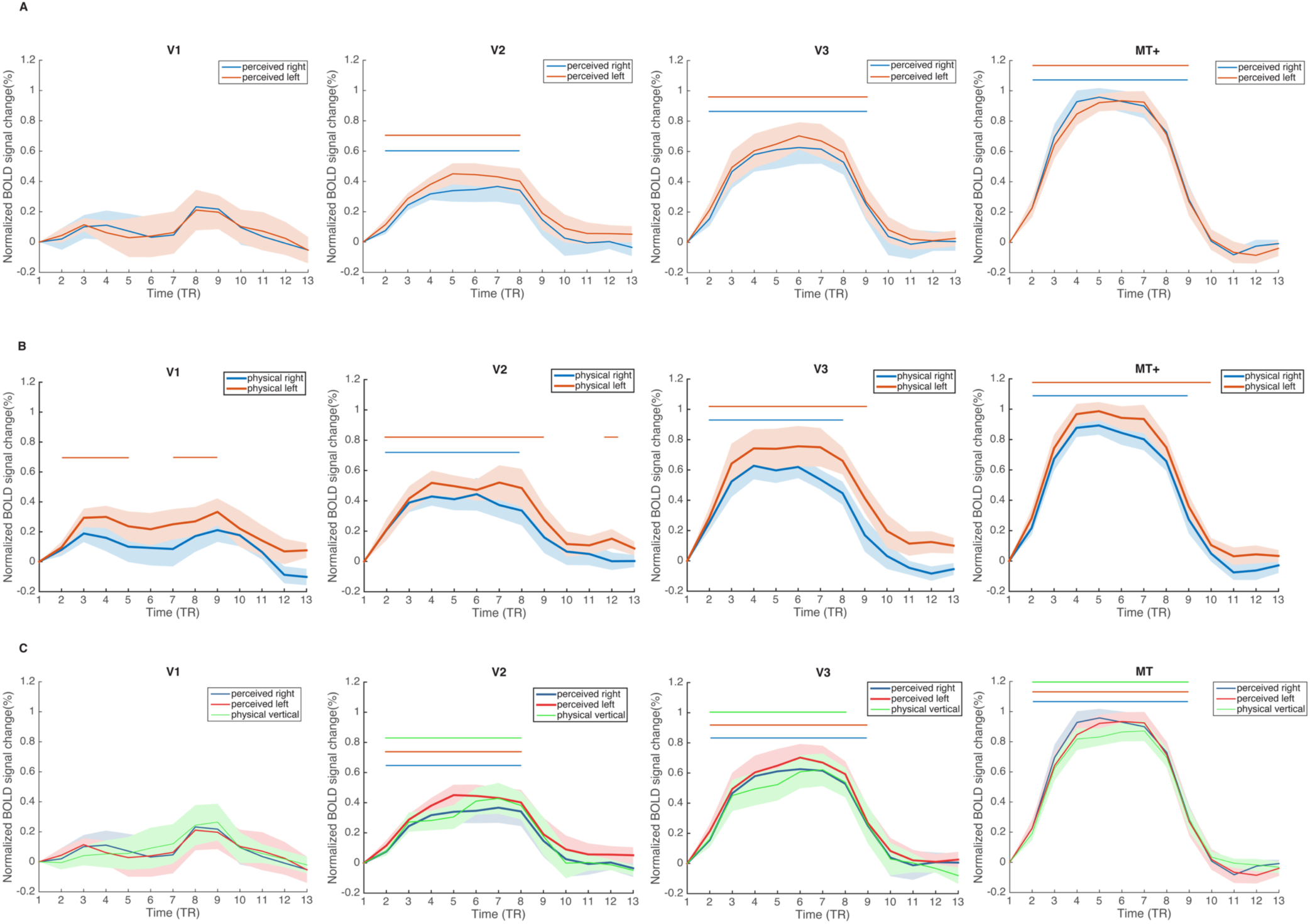
Related to Figure 3. Mean BOLD time course from MPROIs within V1, V2, V3 and MT+. (A) two double-drift stimuli with perceived right vs. left but physically vertical motion directions. (B) two control stimuli with matched physically right vs. left motion direction. (C) two double-drift stimuli with internal motion that drives the perceived paths to appear leftward or rightward and the vertically moving control stimulus with no internal motion. Error bars represents ± 1 SEM, horizontal lines at the top of each figure represent time points with significant above-baseline activity for each stimulus condition (*p* < .05). Paired samples t-tests showed no significant differences between the two double-drift stimulus conditions, the two control stimulus conditions, or between the double-drift stimuli (with internal motion) and control stimulus (with no internal motion) that shared the same physical (i.e. vertical) but different perceived motion direction (i.e. illusory left or right direction) (*p*s > .1).

**Figure S4.**
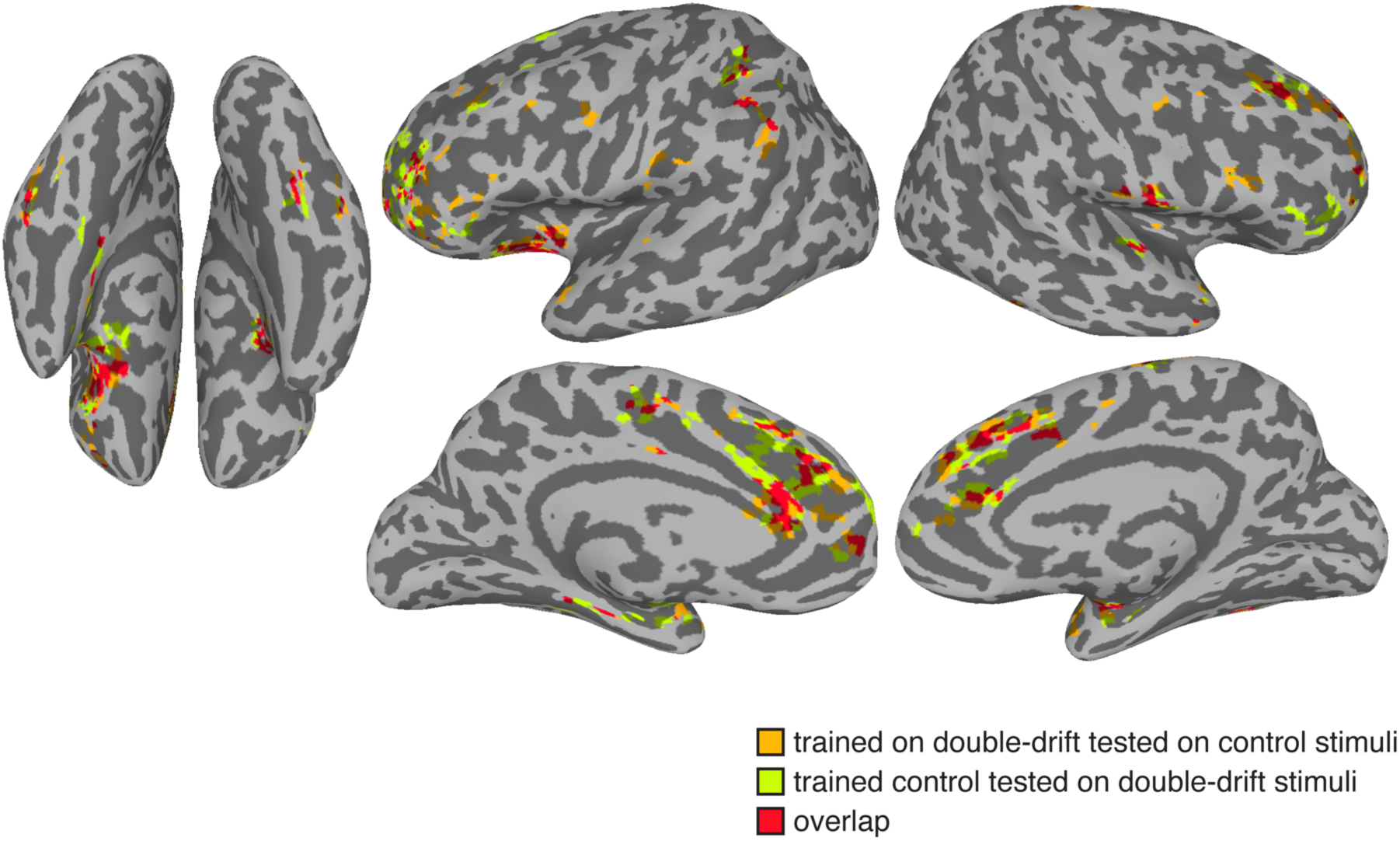
Related to Figure 6. Cluster-thresholded searchlight map with significant above-chance cross-decoding accuracy where the grand mean of each condition was removed within each searchlight during the analysis. Orange represents significant clusters when training the classifier on double-drift and tested on the control stimuli. Green represents significant clusters when training the classifier on the control and tested on the double-drift stimuli. Results were thresholded at *p* = .01 and FDR corrected across clusters at *p* < .05.

**Table S1.**
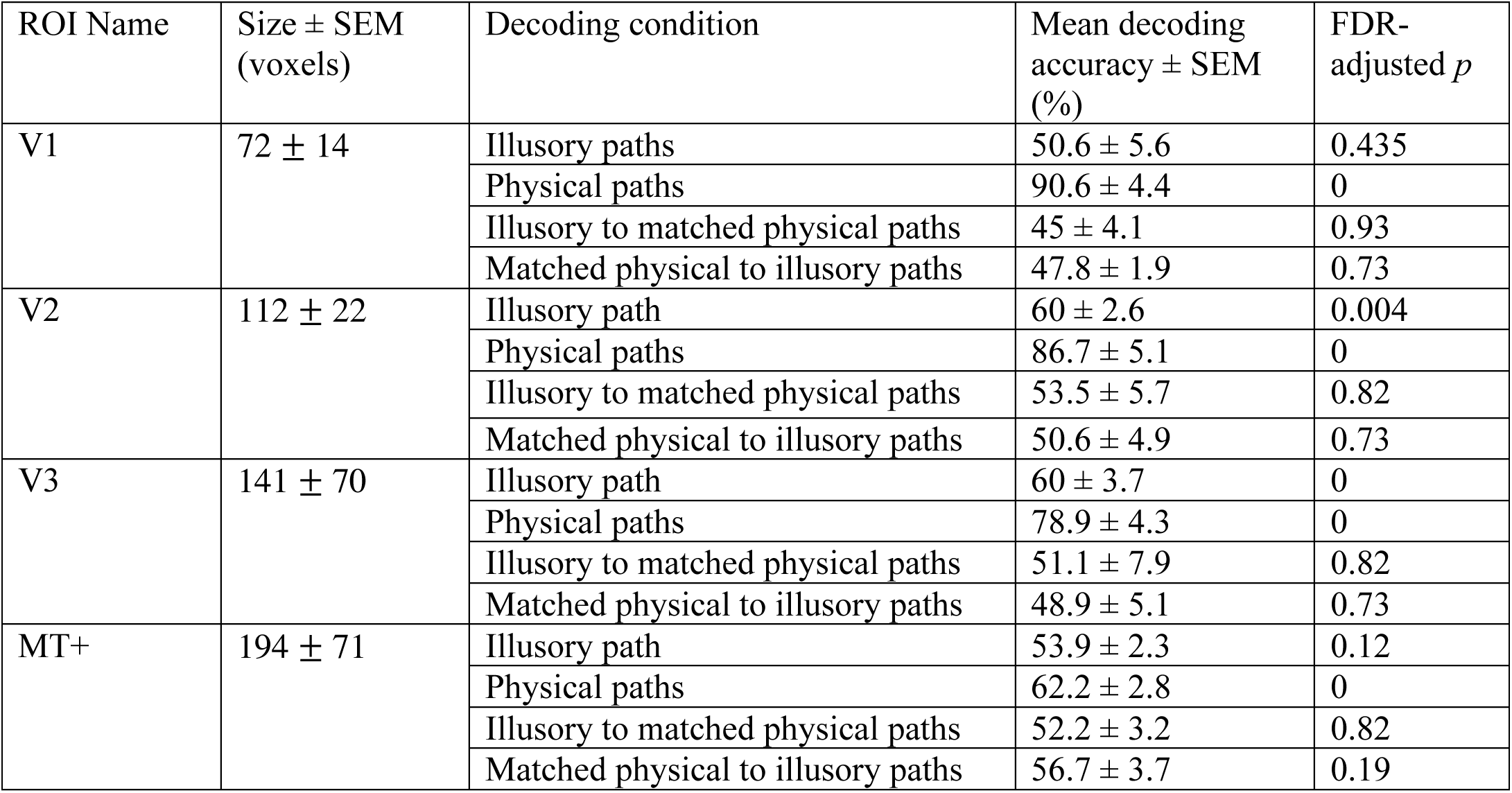
Related to Figure 3. Decoding performances and statistical results in each MPROI.

| ROI Name | Size ± SEM (voxels) | Decoding condition | Mean decoding accuracy ± SEM (%) | FDR-adjusted $p$ |
| --- | --- | --- | --- | --- |
| V1 | 72 ± 14 | Illusory paths | 50.6 ± 5.6 | 0.435 |
| | | Physical paths | 90.6 ± 4.4 | 0 |
| | | Illusory to matched physical paths | 45 ± 4.1 | 0.93 |
| | | Matched physical to illusory paths | 47.8 ± 1.9 | 0.73 |
| V2 | 112 ± 22 | Illusory path | 60 ± 2.6 | 0.004 |
| | | Physical paths | 86.7 ± 5.1 | 0 |
| | | Illusory to matched physical paths | 53.5 ± 5.7 | 0.82 |
| | | Matched physical to illusory paths | 50.6 ± 4.9 | 0.73 |
| V3 | 141 ± 70 | Illusory path | 60 ± 3.7 | 0 |
| | | Physical paths | 78.9 ± 4.3 | 0 |
| | | Illusory to matched physical paths | 51.1 ± 7.9 | 0.82 |
| | | Matched physical to illusory paths | 48.9 ± 5.1 | 0.73 |
| MT+ | 194 ± 71 | Illusory path | 53.9 ± 2.3 | 0.12 |
| | | Physical paths | 62.2 ± 2.8 | 0 |
| | | Illusory to matched physical paths | 52.2 ± 3.2 | 0.82 |
| | | Matched physical to illusory paths | 56.7 ± 3.7 | 0.19 |

**Table S2.** Related to Figure 5. Significant clusters found in the within-condition classification searchlight analysis (thresholded at *p* = 0.01 and FDR-corrected across clusters at *p* < 0.05).

| Decoding condition | Size (voxels) | Peak |  |  |  | areas |
| --- | --- | --- | --- | --- | --- | --- |
|  |  | MNI coordinates |  |  | Classification accuracy (%) |  |
|  |  | X | Y | Z |  |  |
| Illusory path | 4720 | 13.5 | -31.5 | 2.5 | 73.3 | Superior frontal gyrus, medial frontal gyrus, anterior cingulate, right parahippocampal gyrus, right superior temporal gyrus |
|  | 2993 | -10.5 | 58.5 | -12.5 | 72.2 | Middle occipital gyri, right precuneus, right parahippocampal gyrus, right angular gyrus |
|  | 1713 | 22.5 | 10.5 | 29.5 | 72.2 | Left cingulate gyrus, left postcentral gyrus |
|  | 294 | -19.5 | 19.5 | 29.5 | 68.9 | Right cingulate gyrus |
|  | 220 | 34.5 | 1.5 | -21.5 | 71.1 | Left parahippocampal gyrus, left superior temporal gyrus |
| Matched physical path | 21761 | 13.5 | 85.5 | 8.5 | 85.5 | Left middle occipital gyrus, superior and middle temporal gyri, left inferior parietal lobule, superior and middle frontal gyri, precentral gyri, postcentral gyri, paracentral lobule, cingulate gyri |

**Table S3.** Related to Figure 6. Significant clusters found in the cross-decoding searchlight analysis (thresholded at *p* = 0.01 and FDR-corrected across clusters at *p* < 0.05).

| Decoding condition | Size (voxels) | Peak decoding accuracy |  |  |  | areas |
| --- | --- | --- | --- | --- | --- | --- |
|  |  | MNI coordinates |  |  | Classification accuracy (%) |  |
|  |  | X | Y | Z |  |  |
| Trained on double-drift tested on control stimuli | 1670 | 7.5 | -34.5 | 26.5 | 67.2 | right superior frontal gyrus, medial frontal gyri, anterior cingulate |
|  | 441 | 4.5 | 22.5 | 2.5 | 65 | Left thalamus |
|  | 204 | 31.5 | 46.5 | 26.5 | 65.6 | Left inferior parietal lobule |
|  | 168 | -49.5 | -1.5 | 47.5 | 64.4 | Right precentral gyrus, right middle frontal gyrus |
|  | 120 | 43.5 | -1.5 | 35.5 | 63.3 | Left precentral gyrus, left middle frontal gyrus |
|  | 86 | -40.5 | -28.5 | -3.5 | 65 | Right inferior frontal gyrus |
|  | 78 | 37.5 | 46.5 | -24.5 | 64 | Left cerebellum |
|  | 48 | 40.5 | 64.5 | 29.5 | 61.7 | Left angular gyrus |
| Trained on control tested on double-drift stimuli | 603 | -7.5 | -22.5 | 26.5 | 65 | medial frontal gyri, anterior cingulate |
|  | 148 | 34.5 | -49.5 | 23.5 | 62.8 | Left superior frontal gyrus |
|  | 134 | 25.5 | 37.5 | 47.5 | 63.3 | Left postcentral gyrus |
|  | 131 | 7.5 | 28.5 | -3.5 | 63.9 | Left thalamus |
|  | 60 | 22.5 | 31.5 | -21.5 | 61.1 | Left cerebellum |
|  | 50 | -37.5 | -4.5 | -21.5 | 62.8 | Right superior temporal gyrus, right parahippocampal gyrus |
|  | 49 | -61.5 | 10.5 | 20.5 | 61.7 | Right precentral gyrus |
|  | 47 | -43.5 | -37.5 | -3.5 | 61.7 | Right inferior frontal gyrus |
|  | 36 | 22.5 | -1.5 | -12.5 | 61.1 | Left parahippocampal gyrus |
|  | 33 | -13.5 | 19.5 | 59.5 | 61.7 | Right precentral gyrus |
|  | 30 | -4.5 | -52.5 | 23.5 | 60 | Right superior frontal gyrus |

**Table S4.** Related to Figure 6. Significant clusters found in the cross-decoding searchlight analysis where the grand mean of each condition was removed within each searchlight during the analysis (thresholded at *p* = 0.01 and FDR-corrected across clusters at *p* < 0.05).

| Decoding condition | Size (voxels) | Peak decoding accuracy |  |  |  | areas |
| --- | --- | --- | --- | --- | --- | --- |
|  |  | MNI coordinates |  |  | Classification accuracy (%) |  |
|  |  | X | Y | Z |  |  |
| Trained on double-drift tested on control stimuli | 517 | -19.5 | -46.4 | 23.5 | 64.4 | right superior frontal gyrus, right medial frontal gyri |
|  | 402 | 19.5 | -46.5 | -0.5 | 65.6 | Left anterior cingulate, left superior frontal gyrus |
|  | 185 | 31.5 | 40.5 | 35.5 | 64.4 | Left inferior parietal lobule |
|  | 145 | -25.5 | 4.5 | -15.5 | 61.7 | Right parahippocampal gyrus |
|  | 119 | -40.5 | 49.5 | -36.5 | 63.9 | Right cerebellum |
|  | 89 | -10.5 | 13.5 | 65.5 | 64.4 | Right superior frontal gyrus, right medial frontal gyrus |
|  | 82 | 4.5 | 31.5 | 2.5 | 64.4 | Left thalamus |
|  | 75 | 46.5 | 49.5 | -24.5 | 66.1 | Left cerebellum |
|  | 75 | -49.5 | -13.5 | 41.5 | 62.2 | Right middle frontal gyrus |
|  | 46 | -10.5 | -31.5 | 8.5 | 61.7 | Right anterior cingulate |
|  | 45 | -16.5 | -37.5 | 5.5 | 61.7 | Right anterior cingulate, right medial frontal gyrus |
|  | 44 | -61.5 | 10.5 | 20.5 | 61.7 | Right postcentral gyrus |
|  | 34 | 46.5 | -4.5 | 32.5 | 61.7 | Left inferior frontal gyrus, left precentral gyrus |
|  | 31 | 10.5 | 19.5 | -12.5 | 62.2 | Left parahippocample gyrus |
| Trained on control tested on double-drift stimuli | 504 | 34.5 | -52.5 | 11.5 | 66.1 | Left middle frontal gyrus, left superior frontal gyrus |
|  | 487 | -13.5 | -28.5 | 38.5 | 65.6 | Right medial frontal gyrus |
|  | 161 | 13.5 | 25.5 | -12.5 | 63.9 | Left cerebellum |
|  | 121 | 28.5 | 43.5 | 32.5 | 63.9 | Left inferior parietal lobule |
|  | 112 | 22.5 | -19.5 | -9.5 | 63.3 | Left inferior frontal gyrus |
|  | 108 | -49.5 | -34.5 | 11.5 | 63.3 | Right inferior frontal gyrus |
|  | 81 | -25.5 | 1.5 | -15.5 | 64.4 | Right parahippocampal gyrus |
|  | 53 | -22.5 | -52.5 | 14.5 | 62.2 | Right superior frontal gyrus |
|  | 48 | 22.5 | 13.5 | -15.5 | 61.7 | Left parahippocampal gyrus |
|  | 42 | 52.5 | 43.5 | 41.5 | 64.4 | Left inferior parietal lobule |
|  | 41 | -10.5 | 16.5 | 59.5 | 65.6 | Right medial frontal gyrus |
|  | 38 | -58.5 | 7.5 | 17.5 | 63.3 | Right postcentral gyrus |
|  | 33 | 10.5 | -13.5 | 56.5 | 61.7 | Left superior frontal gyrus |
|  | 32 | 7.5 | 10.5 | 44.5 | 61.1 | Left paracentral lobule |
|  | 30 | -31.5 | 37.5 | -18.5 | 61.1 | Right cerebellum |

## KEY RESOURCES TABLE

|  | SOURCE | IDENTIFIER |
| --- | --- | --- |
| Software and Algorithms |  |  |
| MATLAB 2015a | MathWorks [52] | https://www.mathworks.com/ |
| AFNI | [59] | https://afni.nimh.nih.gov/ |
| FreeSurfer | [61] | http://surfer.nmr.mgh.harvard.edu |
| Psychophysics Toolbox | [53] | https://psychtoolbox.com/ |
| R version 3.3.2, RStudio interface | [57,58] | https://www.r-project.com/ |
| PyMVPA | [63] | http://www.pymvpa.org/ |

